# What the odor is not: Estimation by elimination

**DOI:** 10.1101/568626

**Authors:** Vijay Singh, Martin Tchernookov, Vijay Balasubramanian

## Abstract

Olfactory systems use a small number of broadly sensitive receptors to combinatorially encode a vast number of odors. We propose a method of decoding such distributed representations by exploiting a statistical fact: receptors that do not respond to an odor carry more information than receptors that do because they signal the absence of all odorants that bind to them. Thus, it is easier to identify what the odor *is not*, rather than what the odor is. For realistic numbers of receptors, response functions, and odor complexity, this method of elimination turns an underconstrained decoding problem into a solvable one, allowing accurate determination of odorants in a mixture and their concentrations. We construct a neural network realization of our algorithm based on the structure of the olfactory pathway.

## I. INTRODUCTION

The olfactory system enables animals to sense, perceive, and respond to mixtures of volatile molecules carrying messages about the world. There are perhaps 10^4^ or more monomolecular odorants [1–3], far more than the number of receptor types in animals (~ 50 in fly, ~300 in human, ~1000 in rat, mouse and dog [4–7]). The problem of representing high-dimensional chemical space in a low-dimensional response space may be solved by the presence of many receptors that bind to numerous odorants [8–14], leading to a distributed, compressed, and combinatorial representation [12, 15–21] processed by activity in networks of neurons [22, 23]. Some mechanisms for such distributed representation propose that each odorant activates specific subsets of neurons [24– 27]. Other models propose that the olfactory network assigns similar activity patterns to similar odors [28, 29] and classifies odors as activity clusters in an online and supervised manner [30]. Population models suggest that odor identity and intensity could be represented in dynamical response patterns [31, 32], where different odors activate distinct attractor network patterns in a winnerless competition [22, 30, 33], or in transient [34] or oscillatory activity [35]. Finally, population activity could be a low-dimensional projection of odor space [36, 37] evolving in space and time to decorrelate odors [38] to maximally separate sparse representations of similar odors [21].

Here, we focus on a simplified inverse problem: odor composition estimation from time-averaged, combinatorial receptor responses. We thus omit receptor and circuit dynamics important in many olfactory phenomena in animals to concentrate on odor sensing combinatorics (also see [18, 25–29, 39, 40]). We propose that receptors that *do not* respond to an odor carry more information about it than receptors that *do*. This is because silent receptors signal that none of the odorants that could bind to them are present. Most absent odorants can be identified and eliminated with just a few such silent receptors. Thus, it is easier to identify what the mixture *is not*, rather than what the mixture is. For realistic parameters, this elimination turns odor composition estimation from an underdetermined to an overdetermined problem. Then, the remaining odorants can be estimated from active receptor responses.

To be specific, we use realistic competitive binding models of odor encoding by receptors [41–44], and propose schemes to estimate odor composition from such responses. The schemes work over a range of parameters, do not require special constraints on receptor-odorant interactions, and work for systems with few receptors sensing odors of natural complexity. We then develop a neural network inspired by the known structure of the olfactory system to decode odors from receptor responses. We provide performance bounds for these decoders on standard tasks such as detecting presence or absence of odorants in mixtures [45], and discriminating mixtures that differ in some components [46, 47]. These algorithmic schemes are designed with prior knowledge of receptor responses to the complete space of relevant odorants, and hence do not apply as presented to biological olfaction. However, we also construct a version of our neural network without such prior information, albeit at the price of producing a sparse, distributed representation of odors which must be subsequently decoded by a trained classifier.

## II. RESULTS

### A. Identifying odorant presence

Suppose we just seek to identify presence or absence of odor components and not concentrations because the task requires it, or when receptor noise is high. In the latter case, stochastic binding dynamics makes exact binding states hard to predict and receptor activation is determined by noise thresholds. Either way, if the concentration is high and evokes above-threshold receptor activity, the odorant is considered present and the receptor is active. Otherwise, the receptor is deemed inactive and the odorant is absent. Key features of our scheme can be explained in this model. Later, we consider realistic competitive binding (CB) models,= that include continuum odorant concentrations and receptor responses, and then network models which do and do not assume knowledge of the number of odorants and sensing matrix.

Consider mixture of *N*_L_ odorants represented by binary vector 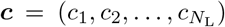, where *c*_*i*_ = 1 represents presence of the *i*’th odorant. Suppose that only *K* odorants are present in the mixture on average. These odorants bind to *N*_R_ receptors whose response is given by the vector 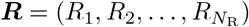. Receptor sensitivity to odorants is given by a matrix *S*, which we assume known. *S*_*ij*_ = 1 indicates that the odorant *j* can bind to receptor *i* and *S*_*ij*_ = 0 means it can not. Suppose that the probability that an odorant binds to a receptor is *s*, i.e., *P* (*S*_*ij*_ = 1) = *s*. Then, on average, each odorant binds to *sN*_R_ receptors and each receptor to *sN*_L_ odorants.

Here receptors respond (*R*_*i*_ = 1) to odors containing at least one odorant binding to them. Without such an odorant, the receptor is inactive (*R*_*i*_ = 0). The receptor thus acts as an ‘OR’ gate, approximating a biophysical model [41–44], with a sigmoidal response function (see below) in situations with high concentration odorants or a sharp threshold and steep response.

Odors encoded in this way can be decoded (estimate ***ĉ***) in two steps (Fig. 1): (**1**) Identify inactive receptors and declare odorants that bind to these receptors as absent, and (**2**) Declare the remaining odorants present. This decoder identifies all odorants in the mixture because, assuming odorants bind to at least one receptor, all receptors that bind to an odorant that is present (*c*_*j*_ = 1) will respond. Hence, its presence will be identified.

**FIG. 1:**
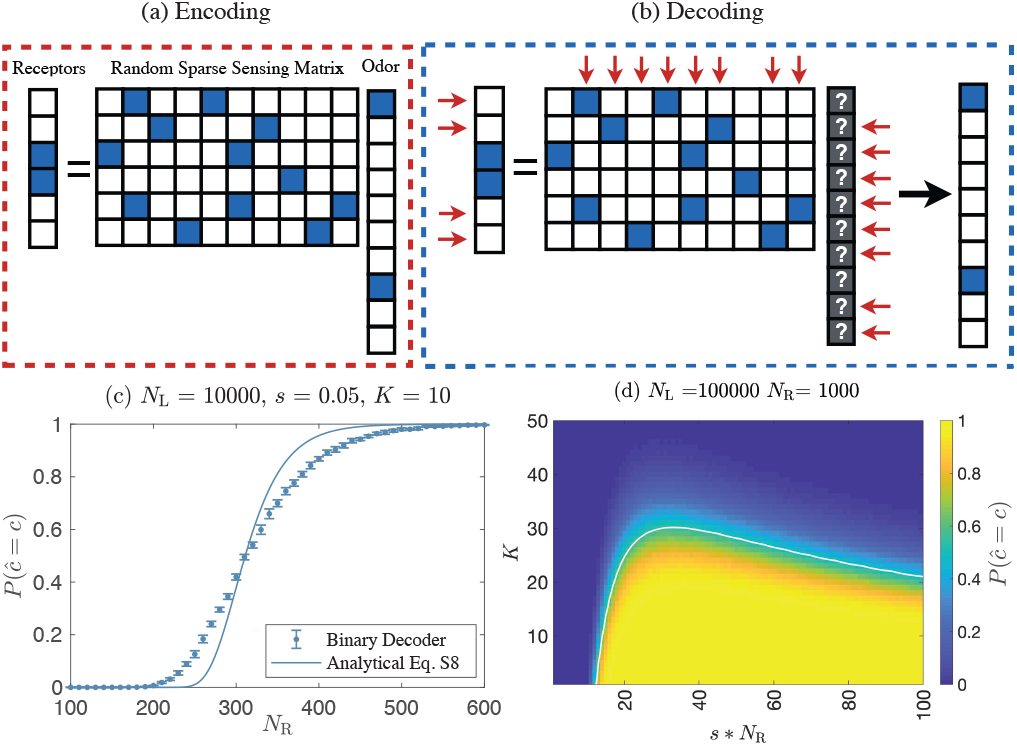
Identifying odorant presence: (**a**) Encoding: The odorant mixture and receptor activity are effectively binary vectors. If an odorant drives receptor activity above threshold, the odorant is considered present and the receptor is said to respond; otherwise the odorant is absent and the receptor is non-responsive. Present odorants and active receptors are indicated as filled elements of corresponding vectors. Here, odorant 1 and 8 are present and receptors 3 and 4 respond. Receptor response is obtained from the sensitivity matrix (filled elements indicate odorant-receptor binding). Here, receptor 1 binds to odorants 2 and 5; receptor 2 to odorants 3 and 7; etc. A receptor is active if the odor contains at least one odorant binding to it; otherwise the receptor is inactive. (**b**) Decoding: (Step 1) Absent odorants are identified from inactive receptors. (Step 2) Remaining odorants are considered present. (**c**) Correct decoding probability (*P* (***ĉ*** = ***c***)) as a function of number of receptors (*N*_R_). Markers = simulations; smooth curve = analytical result (Eq. S8). *P* (***ĉ*** = ***c***) measured as a fraction of correct decodings over 1000 trials with random choices of odor mixture and sensitivity matrix (see SI [48]). Mean and error bar (± 1 standard deviation) computed over 10 replicate simulations (1000 trials each). (**d**) *P* (***ĉ*** = ***c***) as a function of the average number of odorants in mixtures (*K*) and the average number of receptors responding to an odorant (*s* ∗ *N*_R_) (number of receptors (*N*_R_) and odorants (*N*_*L*_) fixed). *P* (***ĉ*** = ***c***) is plotted as a function of *s* ∗ *N*_R_, since *s* and *N*_R_ appear in this combination (Eq. 2 and Eq. S8). *s* ∗ *N*_R_ (average number of receptors binding to an odorant), determines whether the odorant is detectable. *P* (***ĉ*** = ***c***) is measured over 10000 trials with random choices of odor mixture and sensitivity matrix. The white curve is estimated by setting the analytical expression in Eq. 2 to 0.5.

False positives are possible because receptors that bind to an absent odorant could have non-zero response because of other odorants present in the mixture. Thus, the missing odorant will be declared present, giving a false positive. The probability of such false positives is approximately (see SI: Identifying odorant presence [48]):

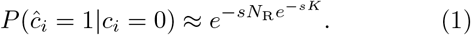

We can derive an approximate probability of correct estimation assuming that each odorant is estimated independently of others (SI: Identifying odorant presence Eq. S8 [48]):

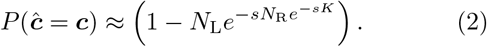

The second term is the approximate probability of false positives with *N*_L_ possible odorants in the environment.

For correct decoding the false positive probability should be low; so, the term in the exponent of Eq. 1 should be large, i.e., *sN*_R_ should be large and *sK* small. This makes sense, as *sN*_R_ is the average number of receptors that an odorant binds. Thus, *sN*_R_ should be large so that many receptors can provide evidence for absence of the odorant by not responding. Also, sufficiently many receptors must be inactive to eliminate all odorants that are absent. For this to happen, the probability that any particular receptor responds to at least one of the K odorants in the mixture should be small. This probability is ≈ *sK* when the likelihood *s* that a given odorant binds to a receptor is small; so we require that *sK <* 1.

The conditions *sN*_R_ *>* 1 and *sK <* 1 are needed because an odorant’s concentration cannot be estimated if it does not bind to any receptor, while converting an under-determined problem into a well determined one requires sufficiently many inactive receptors. Put otherwise, for fixed numbers of receptors and odorants, with fixed odor component complexity *K*, receptor sensitivity should be sufficiently high to ensure coverage of odorants, but small enough to avoid false positives.

These considerations can be combined with the observed sensitivity of olfactory receptors (*s* ~5% for mammals [10]). For typical mammalian parameters ({*N*_L_, *K, N*_R_, *s*} ~ {10^4^, 10, 500, 0.05}), the estimated false positive probability is low (*P* (*ĉ*_*i*_ = 1 | *c*_*i*_ = 0) ~10^*−*7^; Eq. 1) and the correct estimate probability is high (*P* (***ĉ*** = ***c***) ~ 0.998; Eq. 2). These are upper bounds because we considered binary, noiseless signals, while computation in the brain is degraded by noise in sensory and decision circuitry and by circuit constraints, as discussed below. However, our result here shows that in principle, and ignoring noise, odor composition is fully recoverable from the sort of combinatorial codes implemented in the nose.

We estimated our scheme’s accuracy using sensitivity matrices with elements taken non-zero with probability *s*, i.e. (*P* (*S*_*ij*_ *>* 0) = *s*). Fig. 1c shows the correct estimate probability (*P* (***ĉ*** = ***c***)) as a function of the receptor number (*N*_R_) for odors containing on average *K* = 10 odorants drawn from *N*_*L*_ = 10000 possibilities. When the number of receptors is low, the correct decoding probability is zero. As the number of receptors increases, we transition to a region where recovery is good. The transition is sharp and occurs when the number of receptors is much smaller than the number of odorants. Our scheme performs well over a range of odor complexities (*K*) and numbers of responsive receptors (*s* ∗*N*_R_), (Fig. 1d). Our expression for the correct decoding probability (SI Eq. S8 [48]) describes the numerical results well, and gives a good estimate of the transition between poor and good decoding as a function of odor complexity and receptor sensitivity (solid lines in Fig. 1c,d; SI Fig. S1 [48]). We will show that including noise reduces decoding accuracy, matching experiments, but the prediction of a sharp threshold between poor and good performance remains.

### B. Identifying odorant concentrations

When noise is low, or integration times are long, fine gradations of odorant concentrations (*c*_*i*_), receptor sensitivities (*S*_*ij*_), and receptor responses (*R*_*i*_) can be discriminated. The response is then well-described as a Hill function [41–44] because odorant molecules compete to occupy receptor binding sites [44]:

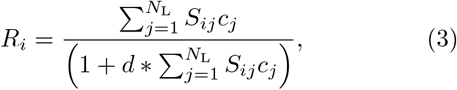

where *c*_*i*_ are odorant concentrations and *d* parameterizes affinity for the receptor. The response is binary when *d* is large (*R* = 0 or 1*/d*), and linear [40, 49–51] when *d* → 0. Synergy, suppression, antagonism, and inhibition, which may be widespread [52], can be included in this model [44]. Below we will first consider a decoder which has explicit knowledge of this receptor response model, and later present neural network models which do not assume knowledge of the number of odorants and response model.

Our decoder can now be modified to estimate both which odorants are present and their concentrations. We start with an under-determined problem because the number of odorants exceeds the number of receptors (*N*_*L*_ *> N*_*R*_). First, we eliminate odorants binding to receptors with below-threshold responses. Thus, an odorant is considered functionally absent if its concentration is low enough that some receptors specific to it respond below threshold. This leaves *Ñ*_*R*_ active receptors responding to *Ñ*_*L*_ candidate odorants. If *Ñ*_L_ ≤ *Ñ*_R_ the problem is now over-determined and can be solved (Fig. 2), even if some absent odorants have not been eliminated. Specifically, we invert the response functions (Eq. 3) relating the *Ñ*_*L*_ odorant concentrations to the *Ñ*_*R*_ responses to get the unknown concentrations.

**FIG. 2:**
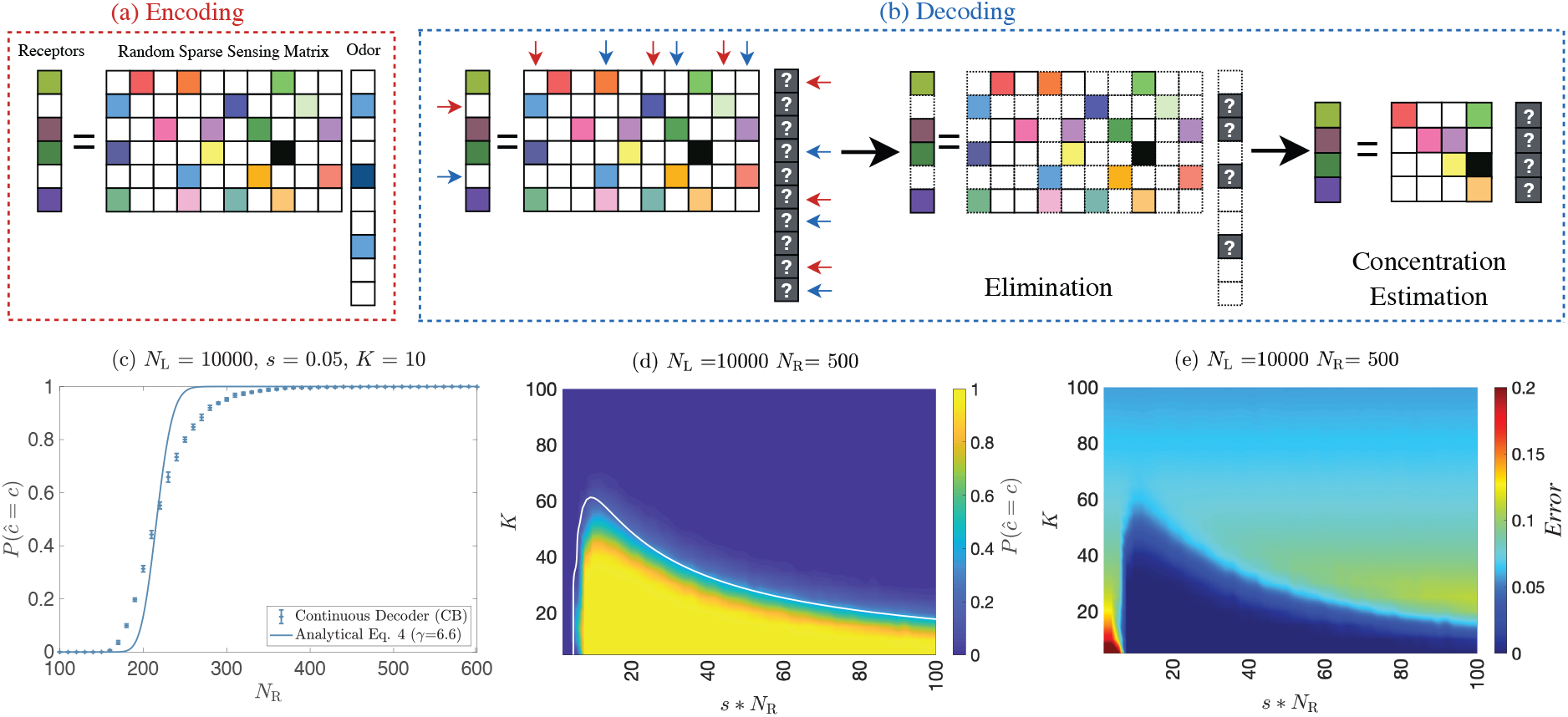
Identifying odorant concentrations: (**a**) Encoding: Odor mixtures are *N*_L_ component vectors with *K* non-zero entries represented by filled elements (color saturation = concentration). Elements of the sensing matrix (rows = receptors, columns = odorants) indicate strength of receptor-odor interactions (color = receptor binding affinity to an odorant; white = zero affinity). Receptor activity = colors, white = inactive. (**b**) Decoding (Step 1 Elimination): Inactive receptors are used to eliminate absent odorants, reducing an under-determined problem to a well-defined one. (Step 2 Estimation): Concentrations of remaining odorants are estimated from responses of active receptors. (**c**) Same as Fig. 1(c), now for the continuous decoder. Mean and error bar (± 1 s.d.) over 10 replicate simulations, each with 1000 trials. The parameter *γ* in the (Eq. 4) was chosen to minimize MSE between the numerical probability and the formula. (**d**) *P* (***ĉ*** = ***c***) as a function of number of odorants (*K*) and *s* ∗ *N*_R_ at fixed *N*_R_. *P* (***ĉ*** = ***c***) calculated over 1000 trials, each with random choices of odor mixture and sensitivity matrix. The white curve is the boundary of the good decoding region, estimated by setting Eq. 4 to 0.5 (*γ* = 3). (**e**) Concentration estimate error (Euclidean distance between actual and estimated values divided by number of odorants; (‖***ĉ***− ***c*** ‖ _2_*/K*)), as a function of number of odorants (*K*) and *s* ∗ *N*_R_ at fixed *N*_R_. Error is low even when recovery is imperfect. Other error measures give similar results (SI Fig. S3: Other measures of estimation error [48]).

Our decoder will eliminate none of the *K* present odorants because all evoke responses. To estimate false positives, let *s* be the probability that a receptor responds to a given odorant (*P* (*S*_*ij*_ *>* 0) = *s*). Then, the number of active receptors is *Ñ*_*R*_ ~ *sKN*_R_ while the number of inactive receptors is about (1 − *sK*)*N*_*R*_. The first inactive receptor eliminates roughly a fraction *s* of the remaining *N*_*L*_ − *K* odorants; the second removes another fraction *s* of the remaining (1−*s*)(*N*_*L*_−*K*) odorants. Summing over these eliminations for all (1 − *sK*)*N*_*R*_ inactive receptors leaves 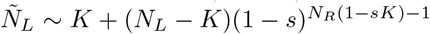 odorants. Typical parameters {*N*_L_, *K, N*_R_, *s*} = {10^4^, 10, 500, 0.05} give *Ñ*_*L*_ ~*K* = 10 which is less than *Ñ*_*R*_ ~*sKN*_R_ = 250; so, in the relevant regime our algorithm leads to an over-determined and hence solvable identification problem.

We can find an approximate analytical expression for the probability of correct estimation (SI: Identifying odorant concentrations [48]) by assuming that the typical number of receptors responding to a mixture exceeds the average odor complexity (*Ñ*_R_ *> K*), while, at the same time, enough receptors are inactive to eliminate absent odorants. This requires *s*(*N*_R_ − *Ñ*_R_) *> γ*, where *γ >* 1 is a parameter depending on the response model (SI: Identifying odorant concentrations [48]). Then,

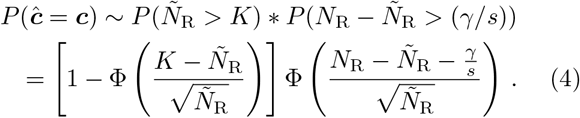

Φ is the normal cumulative distribution function.

To estimate the probability of correct decoding (*P* (***ĉ*** = ***c***)), we generated sparse odor vectors with *K* odorants on average. We drew concentrations from a uniform distribution on the interval [0, 1). Elements of the sensitivity matrix were chosen non-zero with probability *s*; non-zero values were chosen from a log uniform distribution (SI: Numerical Simulations; similar results with other distributions in SI Fig. S2 [48]). With few receptors, the correct decoding probability vanishes. But the probability transitions sharply to finite values at a threshold *N*_R_ much smaller than the number of odorants (Fig. 2c). Odor compositions are recovered well for a range of parameters (Fig. 2d,e), if receptors are sufficiently sensitive *s* ∗ *N*_R_ *>* 6. Odors with the highest complexity are decoded when *s* ∗ *N*_R_ ~ 7 − 15. We quantified the error in odor estimates and found that even when decoding is not perfect there is a large parameter space where the error is small (Fig. 2e). These results have weak dependence on the number of odorants (SI Fig. S4 [48]).

With ~ 300 receptors like human, our model predicts that odors with most components can be decoded with *s* ~ 3 −5%, while with ~ 50 receptors like *Drosophila* we need greater responsivity *s* ≳ 13% for best performance. Interestingly, human receptors have *s* ~ 4% [10] while in the fly *s* ~ 14% [53].

Our decoder can be modified to incorporate other biophysical interactions between odorants and receptors. If non-competitive interactions are known for a receptor-odor pair, the corresponding response model can be used instead of the CB model (Eq. 3). Odorant suppression of receptor responses can be included in the algorithm. If responses are attenuated but not completely silenced, the algorithm goes through as before, albeit with a response model that includes the attenuation. Even if the suppressive interactions are strong, so long as such odor-receptor interactions are sparse, as experiments suggest [9, 10], the algorithm can be modified to include receptor silencing by ignoring odorants and receptors with strong suppressive interactions in the elimination step, and including them in the estimation step. We leave a detailed investigation to future work.

### C. Noise and decision making

To study how noise degrades performance relative to an ideal decoder in our setting, we considered a “cocktailparty problem” where an agent seeks to identify presence or absence of a component in an odor mixture. We then modeled noisy decision-making as a two-step process: (1) Internally representing the mixture using the decoder of odor presence described above, and (2) Using noisy higher level processes to decide on presence or absence of the target odorant based on the output of the estimation step (Fig. 3a).

**FIG. 3:**
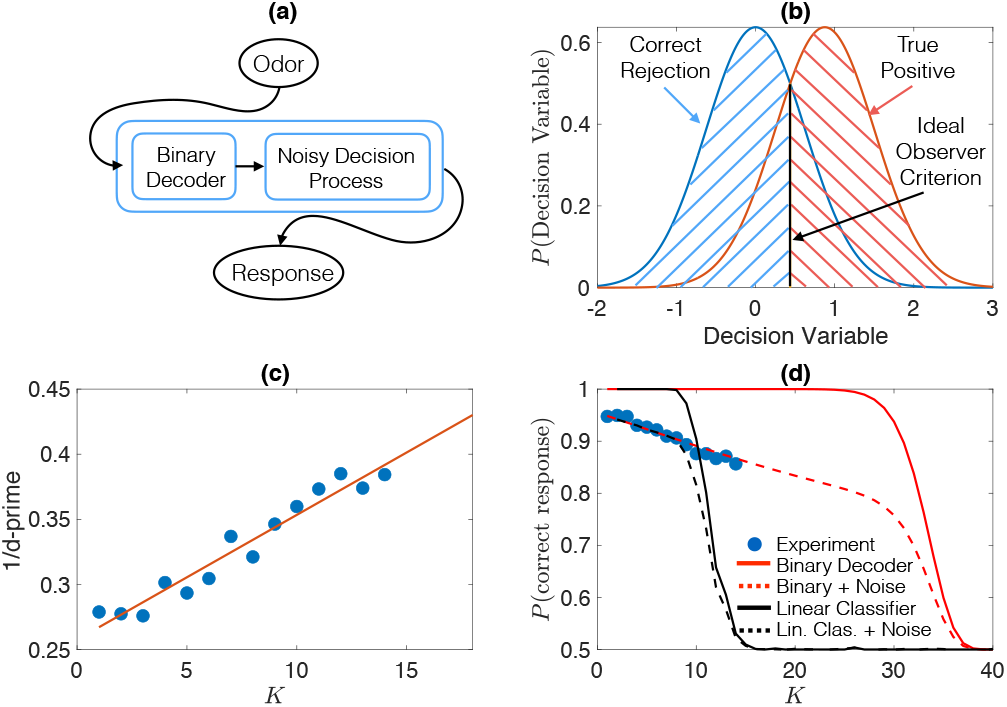
Effect of decision noise: (a) Schematic of a two step noisy decoder. (b) We assume normally distributed decision variables with mean 0 when the target is absent, a mean equal to the probability of correct detection for the odorant presence decoder, *P* (*ĉ* = *c*), when the target is present, and identical standard deviations in both conditions. The ideal observer detection threshold is indicated. Probability of correct response equals probability of correct rejection plus probability of correct detection. (c) 1/d-prime estimated from an olfactory cocktail-party task of detecting odorants in *K*-component mixtures [45] (details in text) follows a linear trend with *K*. (d) Probability of correct detection of presence/absence of an odorant in a *K*-component mixture. Blue markers = fraction of correct responses (true positive + correct rejection). Continuous lines = noiseless prediction for our decoder (red) vs. linear classifier (black). Dashed lines = prediction including noise determined from (c) for our decoder (red) vs. linear classifier (black). Parameters: number of odorants *N*_L_ = 10^4^, number of receptors *N*_R_ = 1000, response sensitivity *s* = 0.05 [10].

We modeled the noisy decision variables derived from activity in a decision circuit [54, 55], by requiring that the baseline-subtracted decision variable should be 0 for absent targets; for targets that are present, the variable should be proportional to the probability *P* (*ĉ* = *c*) = *p* of correct detection. In both cases the decision variable is distributed around the desired value with a standard deviation determined by noise. An ideal observer asks whether the target odorant is more likely to be present or absent, given the observed value of the decision variable and its distribution in the two cases (Fig. 3b).

Next, to derive a realistic noise model, we considered experiments where mice were trained to report a target odorant in mixtures of up to 14 of 16 odorants of identical concentration, any of which could be present or absent [45]. Mice reported presence/absence of all targets with high accuracy (*>* 80%), suggesting that they could learn to identify all mixture components. Decision noise can be directly estimated from this data. Briefly, for Gaussian decision variables (Fig. 3b), we can estimate the standard deviation from the hit rate (fraction of correct detections) and false alarm rate (fraction of incorrect detections). Signal detection theory [56] relates signal to noise ratio (SNR; also called d-prime) of this go/no-go task to the hit/false alarm rates as: SNR = d-prime = z(hit) - z(false alarm), where z is the z-score. This analysis gave the SNR for mice as a function of the mixture complexity *K* ([45]; Fig. 3c). We estimated SNR at other values of *K* by extrapolating the experimental relationship (Fig. 3c, red line). For a Gaussian decision variable with the same standard deviation *σ* in both conditions, and a difference in means of *μ*, standard theory [56] gives SNR = *μ/σ*. We took the noise standard deviation in our model to be *σ* (estimated from the data for each *K*) times a constant chosen to minimize the mean squared difference between theory and experiment.

Since we derived our noise model from data in mice, we constructed a decoder with *N*_*R*_ = 1000 receptors and response sensitivity *s* = 0.05 [10]. Without noise (*σ* = 0), our decoder predicts essentially perfect performance for identification of missing odorants in odors with up to ~ 27 components, and a sharp fall-off thereafter (Fig. 3d, red line). Adding noisy decisions (Fig. 3a,b) leads to the dashed red line in Fig. 3d. Interestingly, there is a good match to the mouse behavioral data in [45] (RMSE between observed and predicted probability of correct estimate = 0.0075). Based on our model, performance in this olfactory cocktail-party problem is predicted to decline linearly as the complexity of odors increases, until there are about 27 odorants. Then, there will be a sharp fall-off in probability of correct detection, approaching chance for odors with ~ 37 components. These predictions depend weakly on the number of odorants (SI Fig. S5 [48]) and strongly on the number of receptors. Thus, if we consider a system with ~ 300 receptors, like human, and assuming similar decision noise, our model predicts performance will be much worse, declining linearly until about 5 − 8 components (*s* = 0.05 − 0.10), then falling sharply to chance at about 14 components (SI Fig. S6 [48]).

We compared these predictions with linear classifiers trained on receptor responses to report whether a target odorant is present. We calculated responses of *N*_R_ = 1000 receptors in the CB model to random *K*-component mixtures drawn from *N*_L_ = 10000 odorants, and trained the classifier on 1000 random mixtures, half containing the target. After training, we estimated classifier performance over 1000 test mixtures, half containing the target, and averaged over 10 random sensitivity matrices, each with different sets of training/test data. This classifier’s performance also declined linearly with odor complexity (Fig. 3d, solid and dashed black lines) but dropped earlier to chance. The RMSE of the linear classifier and the behavioral experiment was 0.1480, higher than our model (0.0075).

### D. Neural network implementation

To implement our algorithm in networks acting on realistic receptors binding stochastically to molecules, we consider a first layer with receptor responses *R*_*i*_ controlled by affinities *S*_*ij*_ between receptors *i* and odorants *j* (concentrations = *c*_*j*_). To mitigate noise we replicate receptors and aggregate activity in a second layer. Reliable non-responses are especially important for us, so we suppress responses by a standard mechanism – recurrent inhibition in this second layer (Fig. 4a) – helping to drive weak, noisy responses below the threshold for activating the next layer of the network. This architecture parallels the olfactory pathway, where each receptor type is individually expressed in thousands of Olfactory Sensory Neurons (Olfactory Receptor Neurons in insects), which are pooled in glomeruli of the olfactory bulb (antennal lobe in insects). The activity of individual mitral cell outputs of each glomerulus (projection neurons in insects) is then suppressed by a widespread inhibitory network of granule cells by an amount that depends on the overall activity of all the receptors [57, 58] (SI Fig. S7 [48]).

**FIG. 4:**
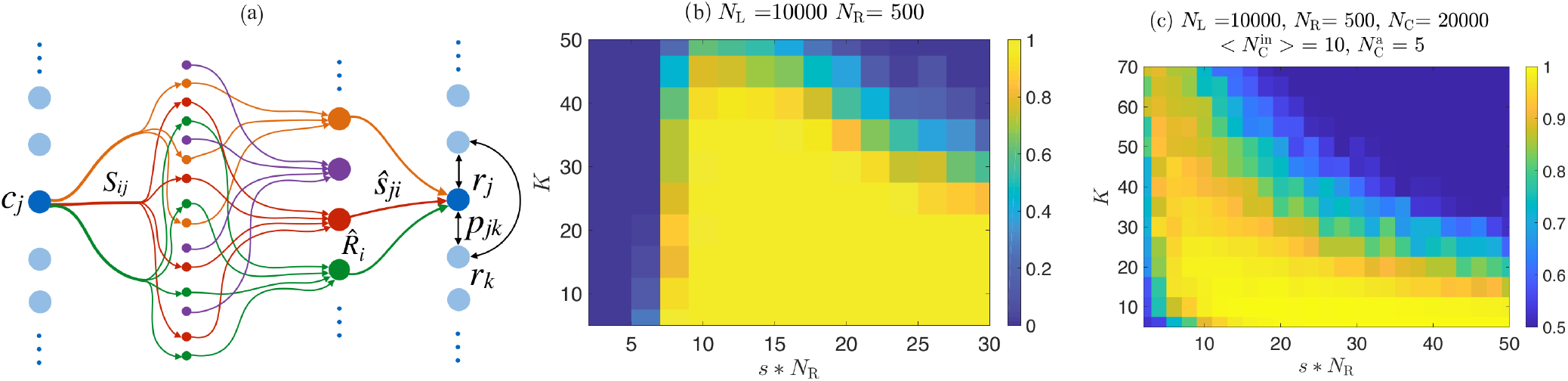
Network decoder: (a) Odorant *c*_*j*_ binds many receptor types (colors). Responses are reliably estimated by averaging multiple receptors of the same type in glomeruli of a second layer where axons of each type converge. Second layer inhibitory interneurons shut down outputs of weakly activated glomeruli. Above-threshold responses are relayed to a readout layer whose units also receive recurrent inhibition from other readout units. Connections for one odorant and readout unit are shown. (b) Probability of correct decoding (*P* (**r** = ***c***)) as a function of odor complexity *K* and *s* ∗ *N*_R_ for *N*_R_ receptors and *N*_*L*_ odorants, with *s* = Probability of odorant-receptor binding. *P* (**r** = ***c***) calculated numerically over 100 trials, with random odor mixtures and sensitivity matrices (SI: Numerical simulations [48]). Correct decoding: Euclidean distance between odor ***c*** and decoded vector **r** is *<* 0.01. (c) Probability of correct classification of presence/absence of a single odorant from the distributed population response in a network with random projection to a third layer with *N*_*C*_ = 20000 units, each of which has an average of 10 inputs 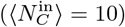, at least half of which must be active (*f* = 0.5) to generate a response. This means that on average the third layer units need at least 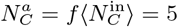 active inputs to produce a response. We used a standard linear classifier, trained with 1000 random odors, half of which contained the target odorant. The classifier was tested on 1000 novel odors. The classifier in panel (c) shows good performance up to higher odor complexities than in panel (b) because it involves a simpler task – i.e. identification of a single odorant, rather than simultaneous identification of all odor components.

We feed second layer outputs 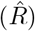 forward with weights *Ŝ*_*ji*_ to *N*_*C*_ third layer units (Fig. 4a). The input to the *j*^th^ unit is gated to implement the elimination step: it is 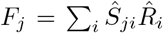 if more than *f* projections to the *j*^th^ unit are non-zero, and vanishes otherwise. This parallels the gating of the projection of the second stage of the olfactory system to Piriform Cortex (Mushroom Body in insects), so that cortical neurons only respond when many inputs are active together [59–62],

The third layer forms a recurrent inhibitory network (weights *p*_*jk*_ *<* 0) with dynamics implementing odor estimation (Fig. 4a). The linearized dynamics of units with non-zero gated input is described by

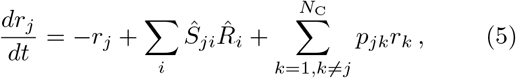

where *r*_*j*_ are responses, and the first term on the right describes activity decay without inputs. *Ŝ* has been restricted to columns and rows associated to active receptors and readout units. Abstractly, the steady state response representing the decoded odor satisfies:

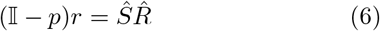

where 𝕀 is the identity; *p* and *Ŝ* are recurrent/feedforward weight matrices; and 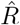 and *r* are response vectors for active receptors and readout units. Thus, the steady state output linearly transforms the gated receptor response.

To illustrate the roles of the feedforward, recurrent and gating structures, suppose the number of readout units and odorants is equal (*N*_*C*_ = *N*_*L*_), sensing is linear with low noise 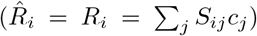, and that gating requires most projections to a responsive unit to be non-zero. Then, at steady state, active units satisfy

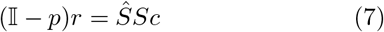

where rows of the odor vector *c* and columns of the sensing matrix *S* have been restricted to present odorants. This readout can directly represent odorants (*r* = *c*) if (𝕀 − *p*)^*−*1^*Ŝ* = *S*^*−*1^, an explicit decoding of the sort considered in [63]. Such an inversion of a rectangular matrix is generally ill-defined because *S* and *Ŝ* will have rank less than the number of concentrations to estimate when there are fewer receptors than odorants. But for sufficiently sparse odors and sensing matrices, we showed that the elimination step removes enough candidates from consideration to give a well defined problem – in our network *S* and *Ŝ* in the active unit dynamics will have sufficient rank to permit the inversion.

Recurrent inhibition allows local solutions to this inversion problem when it is well-defined if we choose

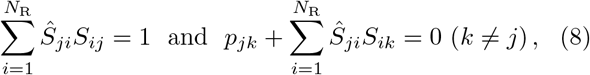

so that (𝕀 − *p*) = *ŜS*. These criteria relate feed-forward weights to the sensing matrix recalling [18, 63], and balance the network unit by unit, compensating feed-forward excitation by recurrent inhibition. Since these constraints relate individual readouts (rows of *Ŝ*) and odorants (columns of *S*), solutions for different pairs can be spliced to construct feedforward and recurrent weight matrices. The balance criterion recalls olfactory cortex where distance-independent projections from pyramidal cells to local inhibitory interneurons produce long-range inhibition [61, 62, 64], and the Mushroom Body in insects where a giant interneuron provides recurrent inhibition. The *N*_L_ + *N*_L_(*N*_L_ 1) equations in (8) are solvable because we have more parameters than constraints: there are ~ *s* ∗ *N*_R_*N*_L_ feed-forward and *N*_L_(*N*_L_ − 1) recurrent parameters in *Ŝ* and *p*. If responses *R*_*i*_ are nonlinear, there will still be enough parameters for decoding, but nonlinear units or multi-layer networks may be needed.

To test the network, we selected sensitivity matrices *S* where odorants bound randomly to a fraction *s* of receptors, and assumed linear responses (Eq. 3 with *d* = 0), representing statistically stable averages over many receptors. We then solved Eq. 8 to find feedforward and recurrent weights (SI: Numerical simulations [48]). Imitating gating of projections to olfactory cortex [59–62], readout units responded if more than 95% of their feed-forward inputs were active. Fig. 4b shows that the network performs similarly to the abstract decoders above.

The architecture above shares features with the olfactory pathway: diffuse but sparse odorant-receptor binding; aggregation and thresholding of noisy responses in the second stage; expansive and strongly gated projections to a recurrent network in the third stage. But the brain does not know the number of odorants or sensing matrix, and so cannot embody networks in which these parameters control the number of neurons or connection weights, at least without learning. However, a variant algorithm works without knowing these parameters. Suppose the feedforward weights to the third layer are sparse, expansive, and statistically random. Strong gating of these projections selects readout units that sample many simultaneously active inputs. Thus, each odorant associates to a sparse readout set, whose activity will be shaped by the dynamics (Eq. 5) to form a distributed odor representation.

This randomly structured network will produce faithful, sparse representations of odor mixtures in the same parameter regime as the elimination-estimation algorithm. To see this, consider a readout population whose activity reflects the presence of odorant *j*. Because of the strong gating, each unit in this population must have a large fraction of its inputs drawn from receptors that respond to *j*. If *j* is absent in a K-component mixture, the probability that a receptor binding *j* remains inactive is ~ *e*^*−sK*^ (SI: Eq. S15 [48]). So, of the roughly *sN*_R_ receptor types that bind to *j*, nearly *sN*_R_*e*^*−sK*^ will be inactive. Taking typical numbers {*K, N*_*R*_, *s*} = {10, 500, 0.05}, ~ 25 receptor types (and the corresponding second stage outputs) will respond to a given odorant, and about 60% of these (~ 15) will be silent if the odorant is absent. The remaining 40% (~10) will respond because of other odorants in the mixture, along with additional receptor types responsive to those odorants. Projecting this activity randomly to the third layer, units in the population representing *j* that sample from silent receptors will be inactive unless sufficiently many new inputs are active because of other odorants in the mixture. This is unlikely because, as discussed above, the strong gating implies that units responding to *j* will have most of their inputs drawn from receptors that do bind to *j*. According to our estimate about 60% of these will be silent if *j* is absent, despite the presence of other odorants in a mixture. Thus the readout unit will not respond. The silenced readout units thus represent *absence* of odorant *j*, which can be explicitly reported by a downstream classifier trained on the sparse third stage activity.

To test this reasoning we constructed a network as described above with statistically random projections to the third layer, and trained a classifier to identify presence of a single odorant based on the third layer population response (Fig. 4c). We found that the classifier showed excellent performance following sparse sensing of odor mixtures with a few tens of components. The classifier in Fig. 4c performs well for odors with higher complexity than in Fig. 4b, because it is performing a simpler task – i.e. detecting a single odorant. Similar classifiers can be built for each odorant of interest, thus forming a classifier layer that explicitly identifies the components of a mixture. A comprehensive future study could also explore, e.g., odor landscapes with different numbers of components, concentration ranges, and statistics; model cortices of different sizes; different statistics and gating in the projections from the second to the third stage; and different kinds of classifiers.

Similar to this decoder, projections from the olfactory bulb to the cortex seem to be statistically random [65] rather than structured, and give rise to a sparse, distributed representation of odors in cortex [17], as opposed to a literal decoding of odorant concentrations. Some authors have proposed that the random projections to cortex are a mechanism for creating sparse, high dimensional representations suitable for downstream linear classification [21, 39, 65], or are evidence for compressive sensing in olfaction [18, 25]. Others have suggested that compressive sensing occurs at the receptors [19, 27], and that the random projections reformat the compressed data for downstream decoding [27]. We propose a complementary view: random projections combine with strong gating to leverage information in silent receptors, enabling network decoding of responses from a small number of receptors.

## III. DISCUSSION

Our central idea is that receptors which do not respond to an odor convey far more information than receptors that do. This is because the olfactory code is combinatorial – each receptor binds to many different odorants and each odorant binds to many receptors. Hence, an inactive receptor indicates that all the odorants that could have bound to it must be absent. Natural odors are mixtures of perhaps 10-40 components drawn from more than 10^4^ volatile molecules in nature [1–3]. If most of these molecules bind to a fraction of the receptors that is neither too small nor too large, odorants that are absent from a mixture can be accurately eliminated from consideration by a system with just a few dozen to a few hundred receptor types. The response of the active receptors can then be used to decode the concentrations of molecules that are present. Our results show that odors of natural complexity can be encoded in, and decoded from, signals of a relatively small number of receptor types each binding to 5-15% of odorants. Perhaps this observation has a bearing on why all animals express ~ 300 receptor types, give or take an O(1) factor, although receptor diversity does increase in larger animals [40] along with the number of neurons in each olfactory structure, the latter scaling with body size [66]. Even at the extremes, the fruitfly and the billion-fold heavier African elephant have 57 ~300*/*6 [4] and 1948 ~ 300 ×6 [67] receptor types respectively.

Our network model, structured similarly to early olfactory pathways, behaves like the abstract algorithms we proposed. Our algorithm and network both show best performance if each of a few dozen to a few hundred receptor types binds to ~5 − 15% of odorants. This is consistent with observations from *Drosophila* to human [10, 53]. Next, our network, like the olfactory system [57, 58], pools receptors of each type into “glomeruli”, and uses lateral inhibition to suppress noise activity. This achieves both reliable responses and non-responses, as required in the elimination step of our algorithm. Our network’s third stage has strongly gated units pooling many glomeruli, most of which must be active to produce responses. The readout units also have large-scale, recurrent, balanced inhibition, like the olfactory cortex [59–62, 64]. Previous work has highlighted that such architectures could enable robust feed-forward odor classification or reconstruction of compressed odor codes [18, 21, 25, 39, 65], and supports both similarity search [28] and novelty detection [29]. We suggest another role for the circuit: to use information in silence to reconstruct odor composition. We also argue that in the absence of information about the dimension of odor space and the sensing matrix, the essential features of our algorithm could be implemented by having random, sparse projections between the second layer and a strongly gated third layer, as seen in the brain [17, 59–62, 65].

The latter perspective involves a subtle point of what it means to “decode an odor. Often we think of decoding as restoration of the “original” signal. We are using the odorant concentrations as the original representation, but could instead think of clouds in molecular shape space, or a points in a space of chemical or biophysical descriptors. Thus, the perspective that odor decoding involves direct recovery of the concentration vector is likely simplistic. Similarly, in vision if a region of the brain decodes the presence of a cat in an image, it does not recover the actual cat, but rather a representation of “catness” that is easy to read. In other words, “decoding essentially involves rewriting information into an easy-to-read format that can be used to generate actions. Thus, the random projections to cortex along with the strong gating (elimination) and recurrent activity (estimation) should be regarded as a population decoding of combinatorial odor information in receptor activity.

We discussed the steady state dynamics of our network decoder, but animal sensing is a highly dynamic affair involving sniffing, active sensing, and transient encounters with odor plumes. In this context, classic work has discussed the role of both oscillatory and transient dynamics in the brain in odor coding and decoding [16, 21–23, 33– 36, 38]. It will be interesting for the future to study how these dynamics interact with the combinatorics of silence that we have discussed, along with the extensive learning and plasticity that occur in the olfactory system.

Future work could also include and study the role of inhibitory and suppressive interactions that have been noted in the nose, where an odorant which does not activate a particular receptor instead suppresses responses of that receptor to other odorants [43, 52, 68]. Odor-invoked inhibitory responses could be formally included in our competitive binding model by including negative entries in the sensing affinity matrix. In this case, the weak response of an ORN may mean that odorants that activate the receptor are absent, or that both excitatory and inhibitory odorants are present at high concentrations. There are potential strategies for including such suppression in our algorithm if odor-receptor interactions are sparse as experiments suggest [9, 10]. Specifically, in the algorithms that assumed knowledge of the receptor response model, we would ignore odorants and receptors with strong suppressive interactions in the elimination step, and include them in the estimation step. If the network decoder does not assume knowledge of the odorant-receptor affinities and instead uses random projections, the readout classifier would have to learn these modifications. We leave detailed analysis for the future.

Our model suggests that estimation and discrimination of complex odors should improve with the size of the receptor repertoire. For example, while humans, dogs and mice might perform similarly at low odor complexity, the latter animals should be better than humans at discriminating more complex odors as they have 2.5 times more receptor types. The quantitative predictions for our model can be determined by studying odor discrimination thresholds as a function of odor complexity for receptor repertoires of different sizes.

Our model also suggests that the information needed for discriminating odor composition may be present in combinatorial receptor representations, contrary to our usual experience of olfaction as a synthetic sense. In fact experiments do show that complex odors differing by just a few components can be discriminated in some circumstances [46, 47]. If the principles underpinning our algorithm are reflected in the brain, as suggested by the analogy with our network model, odors that bind to inactive receptor types should be largely eliminated. We could test this by blocking receptor types pharmacologically, or via optogenetic suppression, and expect that animals will behave as if odorants binding to suppressed receptors are absent, even if other receptors do bind them. Finally, in our model, odors can be decoded well (yellow regions in Figs. 1,2,4) if they have fewer than *K*_max_ components, where *K*_max_ is determined by the number of receptor types (*N*_R_) and the fraction of them that bind to typical odorants (*s*). We can test this by measuring *N*_R_ and *s* for different species and characterizing discrimination performance between odors of complexity bigger and smaller than *K*_max_ (see Fig. 3).

Our algorithm differs from compressed sensing [69–72] which uses a dense, linear sensing matrix to represent high-dimensional sparse vectors in a low dimensional signal, which is decoded through constrained minimization. We similarly assume sparsity of the input vector, but we do not assume linear sensing, and our sensing matrices are sparse, like known odorant-receptor interaction matrices [10, 53]. Also, our decoding mechanism exploits sensory silence, rather than imposing sparsity in the decoded vector [73]. Our approach cannot provide the same general decoding guarantees as compressed sensing, but it succeeds well in the relevant regime of parameters.

Finally, our algorithm may have applications for decoding complex odors detected by chemosensing devices like electric noses [74, 75]. In this engineered setting, the target odorants and response functions are explicitly known so that our method of “Estimation by Elimination” can be precisely implemented.

## Acknowledgements

VS was supported by the University of Pennsylvania Computational Neuroscience Initiative and NIH-SC2GM140945. VB was supported by Simons Foundation MMLS grant 400425, and NSF grants PHY-160761 and PHY-1734030. VB thanks the Kavli IPMU for hospitality. We are grateful to Vikas Bhandawat for useful communications.

## Supplementary Information

### I. ANALYTIC ESTIMATE OF THE PROBABILITY OF CORRECT DECODING

#### A. Identifying odorant presence

We want the probability *P* (***ĉ*** = ***c***) that the decoded vector ***ĉ*** equals the input vector ***c***, i.e., the corresponding elements of the vectors ***ĉ*** and ***c*** are equal. Assuming statistical independence of the decoding of each odorant, we can write

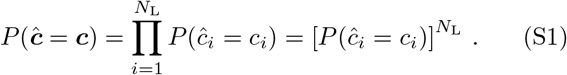

The assumption of independence is an approximation that we will validate by comparing with the full numerical results.

The decoded concentration *ĉ*_*i*_ could be equal to *c*_*i*_, if either both of them equal 1 or both of them equal zero. Thus, the term in the square bracket in Eq. S1 can be written as:

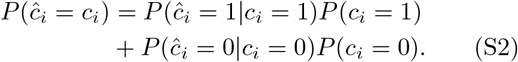

where *P* (*c*_*i*_ = 1) = *K/N*_L_ = *α* is the probability that an odorant is present in the mixture, and *P* (*c*_*i*_ = 0) = (1 − *α*).

The decoder guarantees that if an odorant *c*_*i*_ is present and there is a receptor *R*_*j*_ that is sensitive to it (*S*_*ji*_=1), then the receptor will respond, and the decoded vector will set the corresponding element *ĉ*_*i*_ to 1. If no receptor is sensitive to this odorant (i.e, ∀*j* : *j* ∈ [1, *N*_R_], *S*_*ji*_ = 0), the decoded element will still be set to 1 by default. So, *P* (*ĉ*_*i*_ = 1|*c*_*i*_ = 1) = 1.

To calculate *P* (*ĉ*_*i*_ = 0 |*c*_*i*_ = 0), recall that in our decoding scheme, *ĉ*_*i*_ = 0 if there exists at least one receptor such that *R*_*j*_ = 0 for which *S*_*ji*_ = 1. Thus,

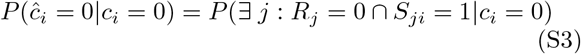

where ∩ is the binary AND operation. The probability on the right is 1 minus the probability that for all receptors either *R*_*j*_ = 1 or *R*_*j*_ = 0 ∩ *S*_*ji*_ = 0. So,

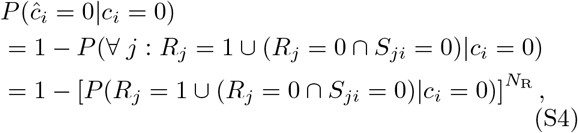

where in the second step we have again made the assumption that the receptors are independent conditional on the response of *c*_*i*_. The quantity in the bracket in Eq. S4 can be written as:

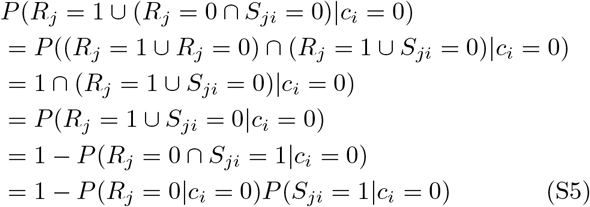

Now, *P* (*S*_*ji*_ = 1 | *c*_*i*_ = 0) = *P* (*S*_*ji*_ = 1) = *s*, where entries of the sensing matrix are chosen to be non-zero independently and with probability *s*.

To calculate *P* (*R*_*j*_ = 0 | *c*_*i*_ = 0) recall that the receptors are OR gates with inputs *S*_*jk*_*c*_*k*_. Thus, for *R*_*j*_ = 0 all terms *S*_*jk*_*c*_*k*_ should be zero. The probability that any one such term is zero is (1 −*sα*). Since we already have *c*_*i*_ = 0, there are (*N*_L_ − 1) additional terms that need to be zero. Hence,

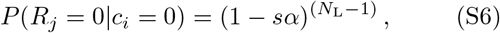

and

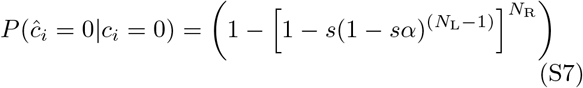

Putting this all together (using Eq. S7 in Eq. S2), we get:

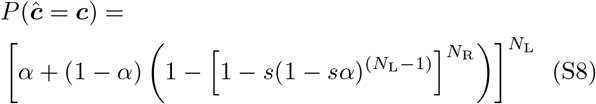

Using Eq. S7, we can also get the (approximate) probability of a false detection as *P* (*ĉ*_*i*_ = 1|*c*_*i*_ = 0) = 1−*P* (*ĉ*_*i*_ = 0|*c*_*i*_ = 0):

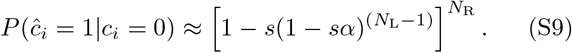

This expression is approximate due to our independence assumptions.

##### 1. Approximation

Since the average number of odorants present in the mixture (*K* = *αN*_L_) is small compared to *N*_L_ and *N*_L_ ≫ 1, we can approximate:

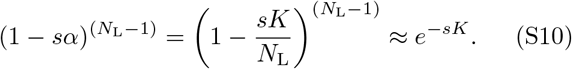

Now, since the odor sensitivity (*s*) is small, so that *se*^*−sK*^ is also small, while *N*_R_ ≫1, we further approximate

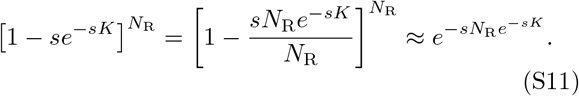

This results in:

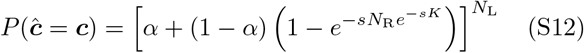

which simplifies to:

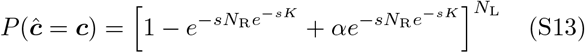

This expression approximates to Eq. 2 in the main text:

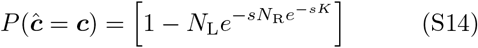

Similarly, Eq. S6 approximates to

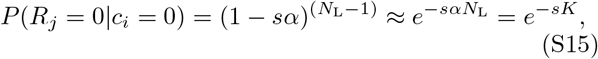

and Eq. S9 approximates to

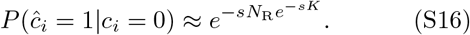

### B. Identifying odorant concentrations

For the continuous decoder to give a unique solution, the number of receptors that respond to the mixture should be larger than the number of odorants with non-zero concentrations (*K* = *αN*_L_). This ensures that the system of equations is over-determined and can, in principle, be solved.

Additionally, the number of receptors that do not respond should be such that the absent odorants can be set to zero. Since every receptor binds to *sN*_L_ odorants on average, we need at least 1*/s* receptors to cover all the odorants. In general, as the entries of the sensitivity matrix are statistically distributed, the number of receptors that do not respond should be larger than *γ/s* for correct odor estimation, where *γ* is a small number greater than 1.

Putting this all together, if *P* (*Ñ*_R_) is the probability of the number of receptors with non-zero response, we are interested in the probability that *P* (*Ñ*_R_ *> K* = *αN*_L_) ∗ *P* (*N*_R_ − *Ñ*_R_ *>* (*γ/s*)). The probability that a receptor responds is:

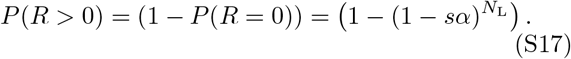

Taking the number of receptors that respond to be a Poisson variable with rate ⟨*Ñ*_R_ ⟩= *N*_R_ ∗*P* (*R>* 0), we can estimate the typical number of receptors that respond. For biologically appropriate parameters {*N*_L_, *N*_R_, *K, s*} ~ {10^4^, 500, 10, 0.05}, the mean number of receptors that respond is ⟨*Ñ*_R_ ⟩~ 200. The standard deviation is 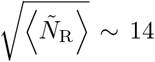 For these values of the mean and variance, we can approximate the Poisson distribution with a Gaussian 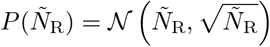. Thus,

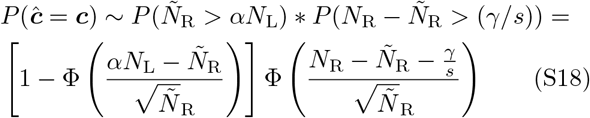

where Φ is the cumulative distribution function of the standard normal distribution.

**FIG. S1:**
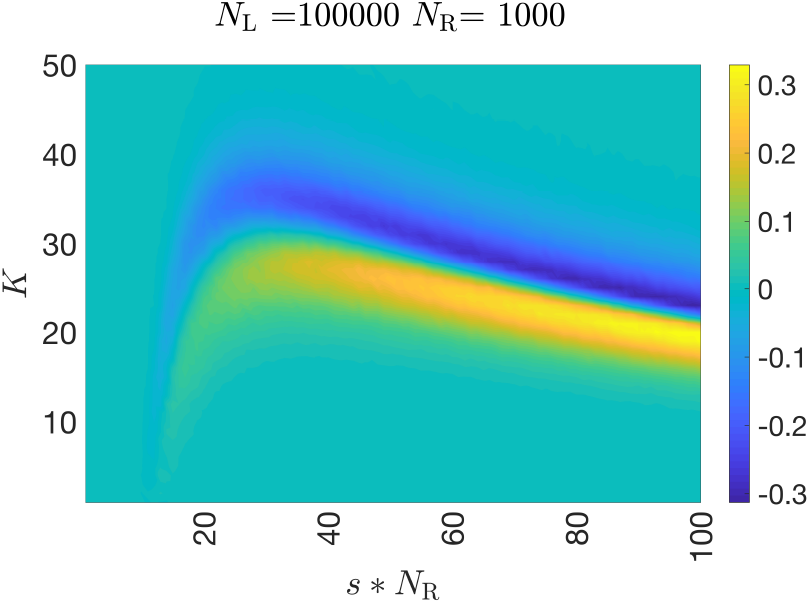
Plot of the difference between *P* (***ĉ*** = ***c***) as given by Eq. S8 and as estimated numerically for the binary decoder. For most parameters the analytical results match the simulations.

### C. Numerical Simulations

#### 1. Identifying odorant presence

For the binary case, the elements of the odor vector were chosen to be non-zero with probability *P* (*c*_*i*_ *>* 0) = *K/N*_L_. The entries of the sensitivity matrix *S*_*ij*_ were chosen to be non-zero with a probability *s*, (*P* (*S*_*ij*_) = *s*). The receptor response was calculated using the binary ‘OR’ function. The decoded concentration *ĉ* was estimated using the two steps described in the main paper. First, the decoded concentration of any odorant to which an inactive receptor is sensitive, was set to zero. All remaining concentrations were set to 1.

#### 2. Identifying odorant concentrations

For the continuous case, the elements of the odor vector were chosen to be non-zero with probability *P* (*c*_*i*_ *>* 0) = *K/N*_L_, and the elements of the sensitivity matrix were chosen to be non-zero with probability *s*, (*P* (*S*_*ij*_ *>* 0) = *s*). The values of the non-zero elements in the odor vector were chosen from a uniform distribution on the interval [0, 1), and for the sensitivity matrix from a log-uniform distribution between 10^*−*1^ and 10^1^. The activity of each receptor was determined using Eq. 3 of the main text (d = 1).

**FIG. S2:**
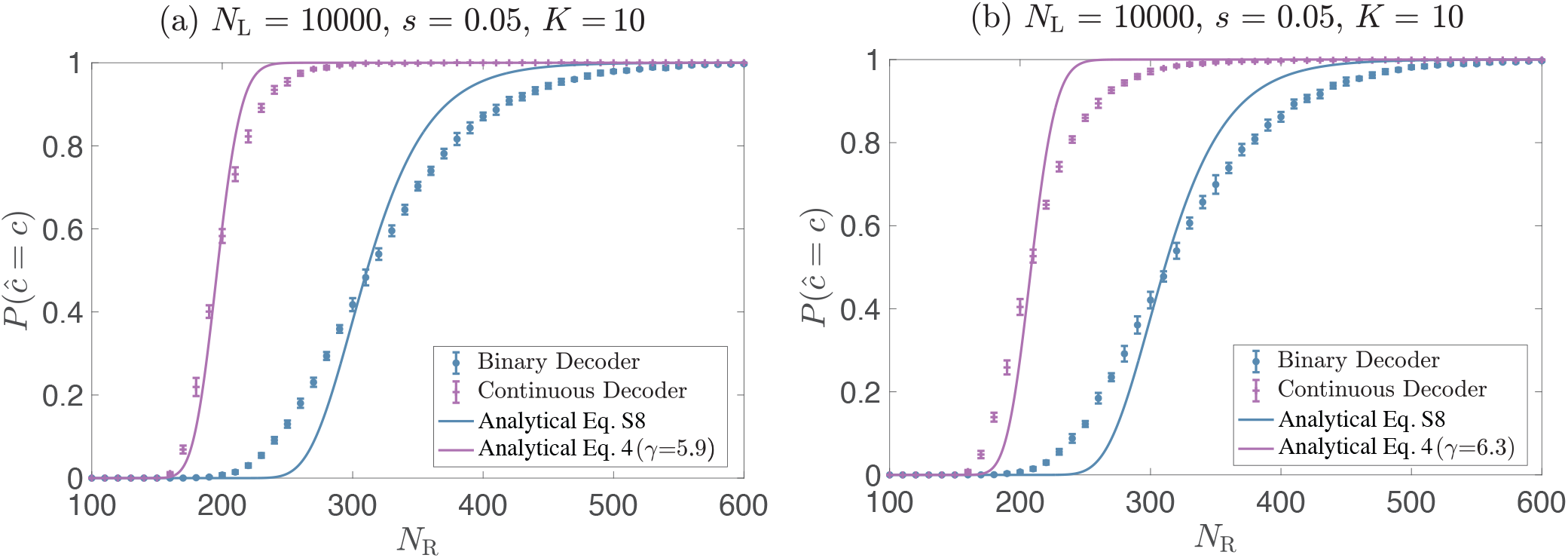
*P* (***ĉ*** = ***c***) **as a function of** *N*_R_ **for the continuous decoder and alternative choices of the sensitivity matrix**. Results for binary encoding are the same as in Fig. 1c and are plotted here for comparison. (a) Uniform distribution: Similar to Fig. 2c except that the non-zero elements of the sensitivity matrix were chosen uniformly at random between [0,1]. (b) Log-normal distribution: Similar to Fig. 2c except that the non-zero elements of the sensitivity matrix were chosen at random from a log normal distribution with the corresponding normal distribution having mean zero and standard deviation 1.

**FIG. S3:**
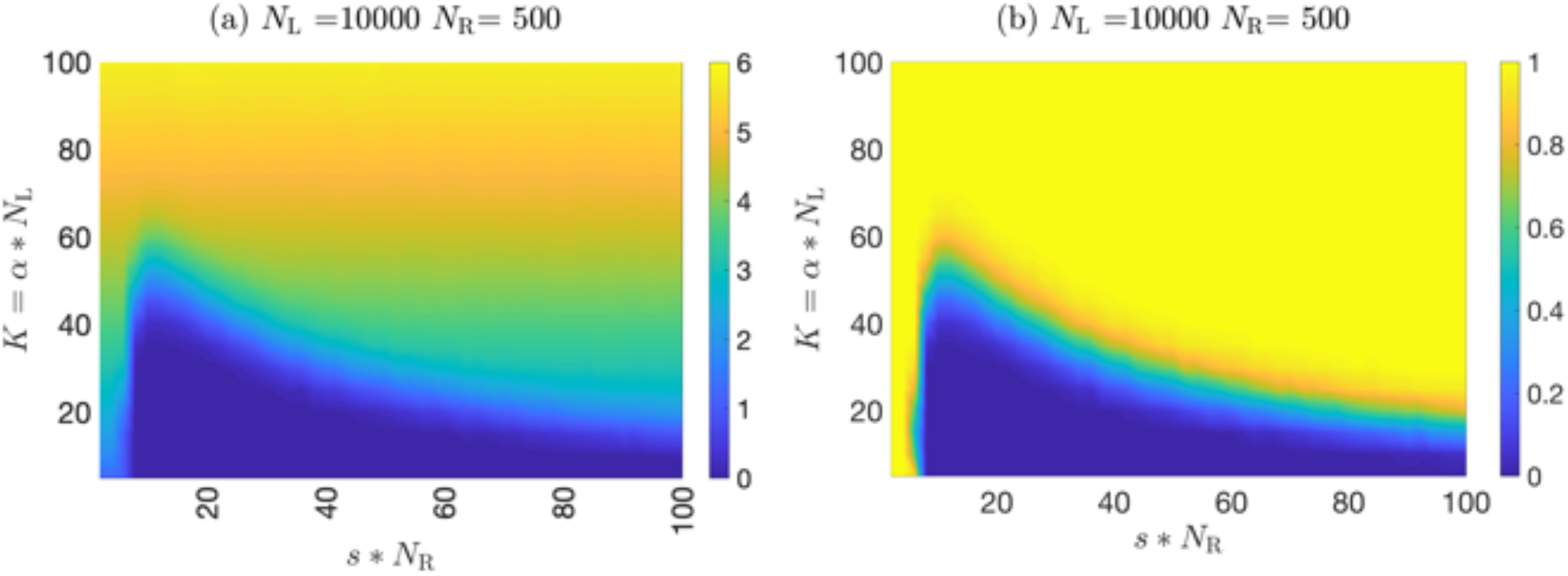
Other measures of estimation error: (a) Total error: *L*_2_ norm (or the root mean square) of the difference between actual and estimated concentrations for the continuous decoder with compressive binding encoding model. (b) *L*_2_ norm of the difference between actual and estimated concentrations divided by the *L*_2_ norm of the actual concentration.

The concentration of any odorant to which an inactive receptor is sensitive was set to zero. After this elimination, let 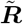 be the vector representing the response of the set of active receptors, 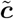 be the vector representing the concentration of the odorants that have not been set to zero, and 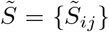 be the *Ñ*_R_ × *Ñ*_L_ sensitivity submatrix over active receptors 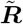 and the remaining odorants 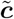. Then, if *Ñ*_R_ *< Ñ*_L_ (non-invertible case), all decoded concentrations were set to zero. Otherwise, the decoded concentrations were given by the vector that minimized the *L*_2_ distance 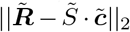. The Levenberg-Marquardt solver with geodesic acceleration from the *GNU GSL* library was used to find the minimum.

Multiple trials were run for each choice of parameters. At the end of each trial, the *L*_2_ norm of the difference between actual and decoded concentration vectors was reported. The trial was considered a success if this norm was less than a threshold of 0.01.

**FIG. S4:**
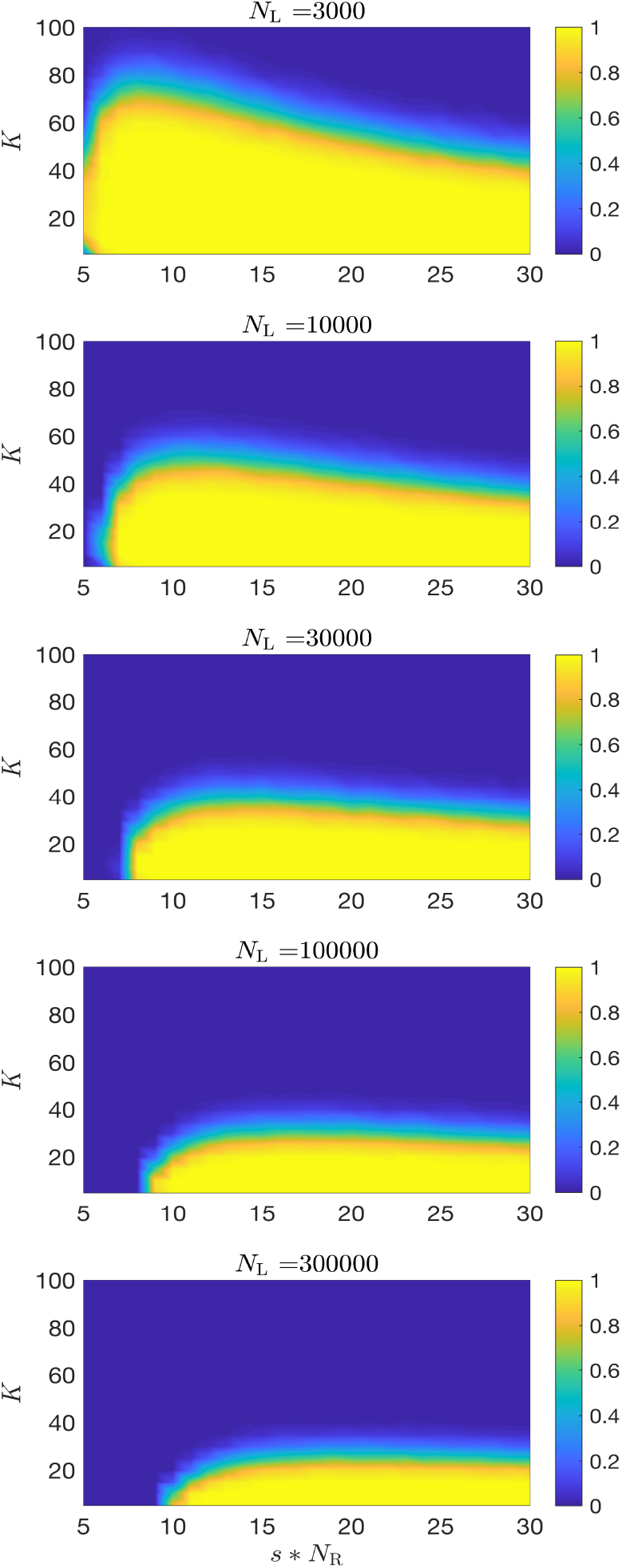
Dependence of *P* (***ĉ*** = ***c***) **on** *N*_L_: *P* (***ĉ*** = ***c***) plotted as a function of odor complexity *K* and *sN*_R_ at a fixed value of number of receptors *N*_R_ = 500. Each panel gives *P* (***ĉ*** = ***c***) for a different value of the total number of possible odorants *N*_L_. The minimum value of *sN*_*R*_ for successful decoding and the optimal value where the most complex odors can be decoded are both relatively independent of *N*_L_.

Simulations were performed in C++. The sensitivity matrix *S* and the odorant concentrations were generated from streams of (pseudo)random numbers drawn by the *Xoroshiro128+* random number generator. Each stream is seeded with a 2^64^ forward jump from the seed of the previous stream. The first stream is seeded from the output of the *SplitMix64* generator initialized by current system time. Random number production as well as vector operation code were optimized using *Intel* ‘s SIMD instruction set.

#### 3. Neural network

To simulate the neural network, we generated random sparse odor vectors and sensitivity matrices. The elements of the odor vector were chosen to be non-zero with probability *P* (*c*_*i*_ *>* 0) = *K/N*_L_, and the elements of the sensitivity matrix were chosen to be non-zero with probability *P* (*S*_*ij*_) *>* 0 = *s*. The value of the non-zero elements were chosen from a uniform distribution on the interval [0, 1). The receptor response was calculated using a linear response model (d = 0 in Eq. 3). To get the feed forward connections *Ŝ*_*ji*_, we first made a matrix 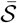 defined as: 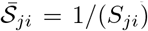 if *S*_*ji*_ is non-zero, and 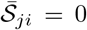 otherwise. The matrix *Ŝ* was then chosen as: 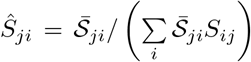. The elements of the recurrent connectivity matrix were obtained as 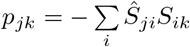.

If more than 5% of the receptors connected to a read-out unit *r*_*j*_ were inactive, the decoded concentration *r*_*j*_ was set to zero. The feed-forward input to the remaining readout units were calculated as 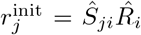. The remaining concentrations were computed as 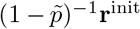, where **r**^init^ is the vector representing the total feed-forward input to neurons that have more than 95% of their receptors active, and 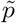 represents the sub-matrix of connection weights between these neurons.

#### 4. Network Classifier

To simulate the network classifier, we generated random sparse odor vectors and sensitivity matrices similarly to the neural network described above. The receptor/glomerular responses were calculated using the competitive binding model. The receptor/glomerular response was projected randomly to a third layer (cortical layer with *N*_C_ neurons): the connectivity matrix was generated so that each second layer unit had a small nonzero probability of connection to each third layer unit such that third layer units received an average of 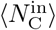 inputs. The connection strengths were sampled from a uniform distribution. A unit in the third layer produced a response if more than a fraction *f* of its inputs was active. If active, the response was calculated using a saturating activation function of the form given in (Eq. 3). For the simulations in the main text, we chose 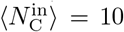 and *f* = 0.5. Thus, for a neuron in the third layer to respond, on average at least 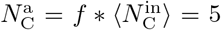 of the inputs needed to be active. We built a linear classifier based on the response of the neurons in the third layer, using the Matlab function *fitclinear*.

To train the classifier, we generated 1000 random sparse odor vectors, half of them containing a specific odorant and the other half not. We calculated the response of the neurons in the third layer for these random odors. We then tested the performance of the classifier on a test set of 1000 different random sparse odors, half of which contained the specific odorant. We calculated fraction of times the classifier correctly identified the odorant to be present or absent.

We repeated this process 10 times, each with a new sample of the odor-receptor sensitivity matrix and receptor-cortex projection matrix. We estimated classifier performance as the fraction of correct identification averaged over these 10 simulations. For the plot presented in Figure 4c of the main text, we repeated the above process for each combination of the parameters *K* and *s* ∗ *N*_R_ with new samples of sensitivity and projection matrices.

**FIG. S5:**
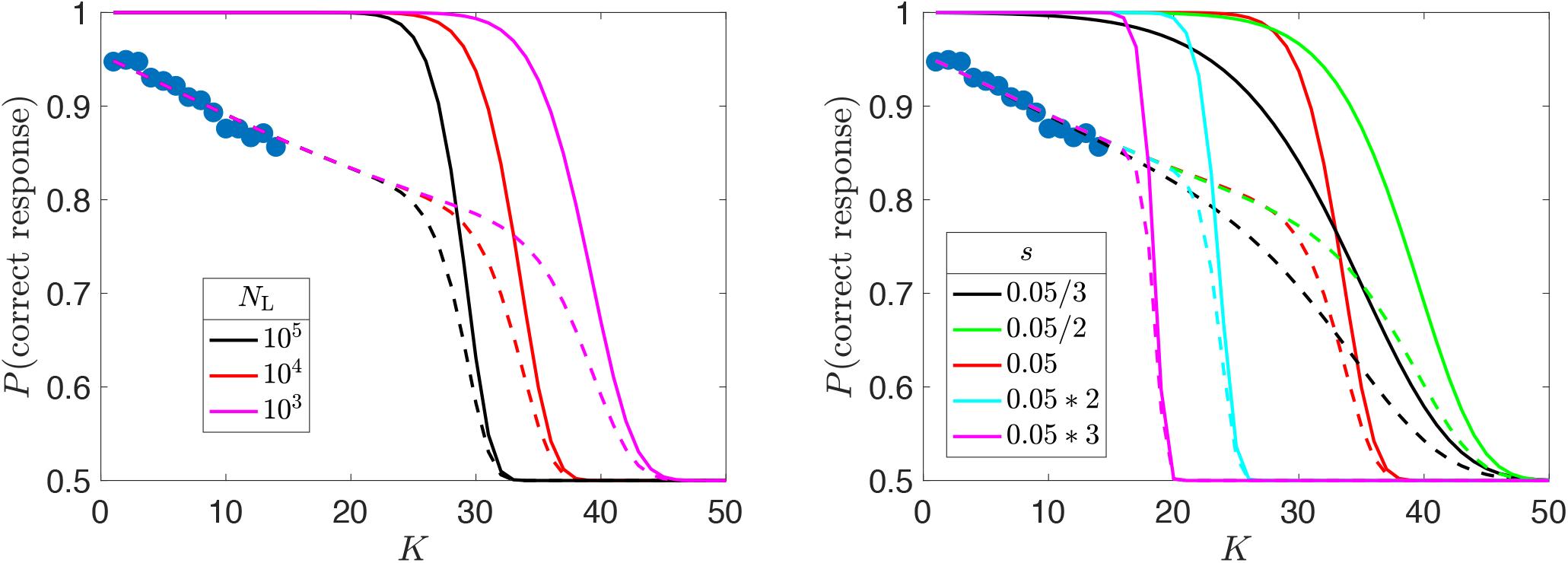
Effect of the number of odorants *N*_L_ and receptor sensitivity *s* on performance in an olfactory cocktail party problem: Probability of correct detection of presence or absence of an odorant in a *K*-component mixture. Compare to Figure 3d. (a) *N*_R_ = 10^3^,*s* = 0.05. (b) *N*_R_ = 10^3^, *N*_L_ = 10^4^.

**FIG. S6:**
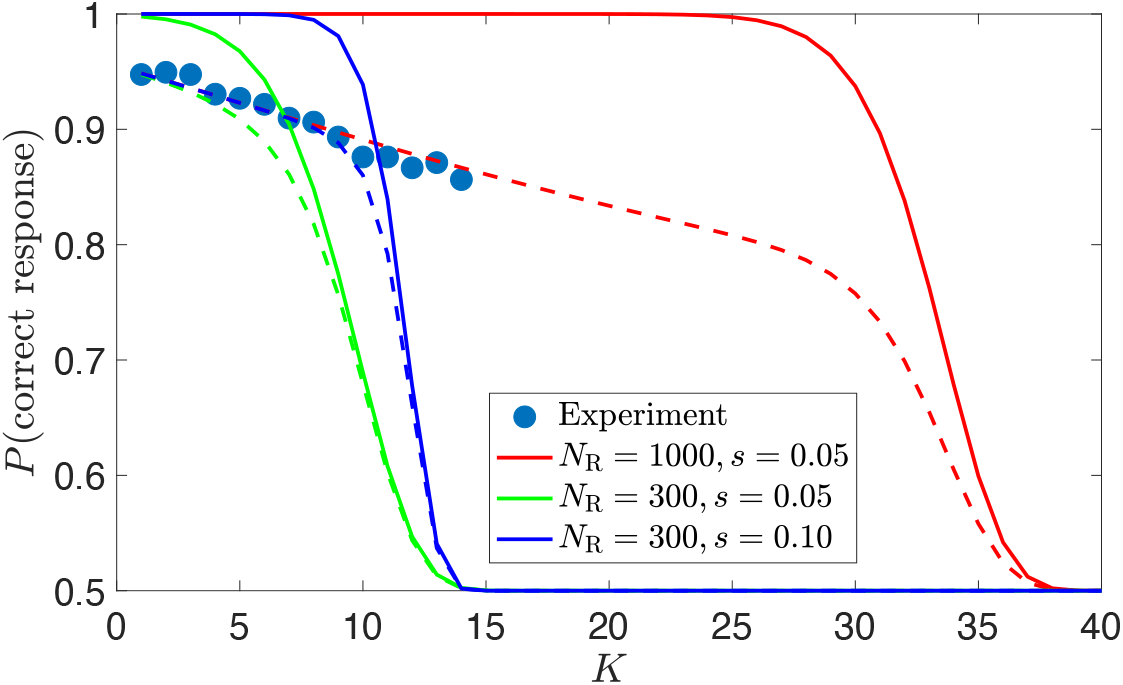
Effect of number of receptors on performance in an olfactory cocktail party problem: Probability of correct detection of presence or absence of an odorant in a *K*-component mixture. Blue markers = fraction of correct responses (true positive + correct rejection) by mice. Continuous lines = prediction in the absence of noise for the binary decoder. Dashed lines = prediction including noise determined from the data in [45] (Fig. 3c main text). Performance with *N*_R_ ~ 300 receptors, like human, is predicted to be lower than with *N*_R_ ~ 1000 receptors, like mouse, falling to chance level at *K* ~ 14. We have shown the prediction with 300 receptors at two values of the receptor-odor sensitivity (*s* = 0.05 (green lines) and *s* = 0.10 (blue lines)).

**FIG. S7:**
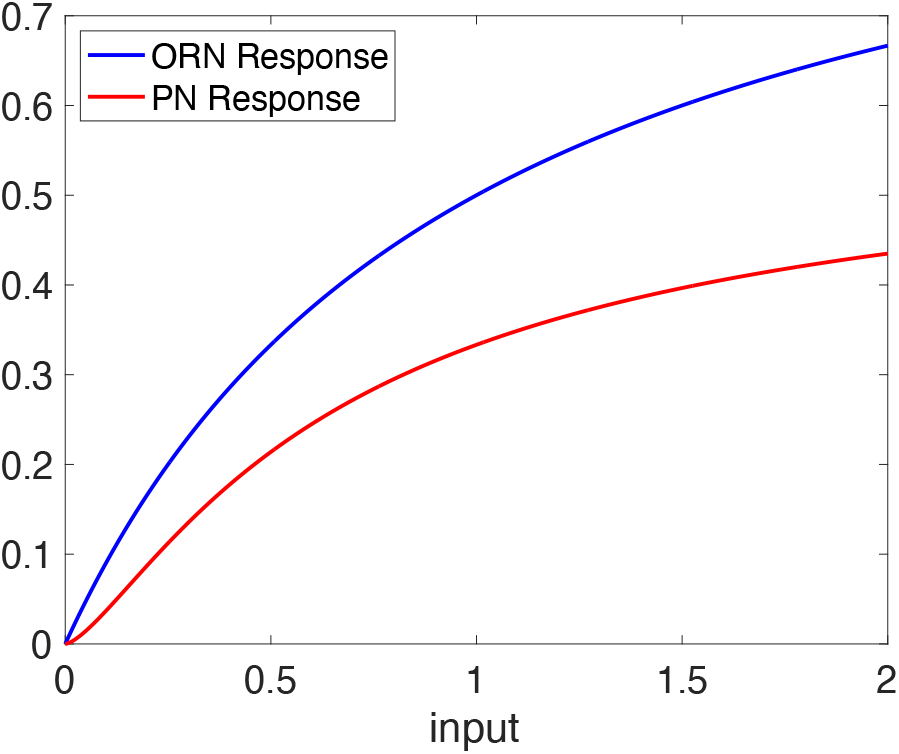
Effect of divisive normalization on output neurons of Antennal Lobe (analog of Olfactory Bulb in insects): The response of the output Projection Neurons (PN) of the Antennal Lobe undergoes a divisive normalization driven by lateral inhibition [56]. The PN output is related to the Olfactory Receptor Neuron (ORN) activity by 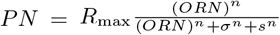, where *σ* parametrizes spontaneous activity, *s* increases in proportion to the overall activity of ORNs, and the coefficient *n* = 1.5. *R*_max_ is a scale determined by the amount of ORN pooling in the glomeruli of the Antennal Lobe. Here we set *R*_max_ = 1 because the overall scale is not relevant to our considerations; rather we are interested in the relative suppression due to overall activity of receptors. Since *n >* 1, the response of each projection neurons is suppressed by the overall activity. While all PNs are suppressed, weakly active glomeruli will be driven closer to, or below, activation thresholds for the next stage of processing. In this figure, the ORN response is given by the competitive binding model, and we chose *σ* = 1, *s* = 1 in arbitrary units to illustrate the suppression. Note that in [56] the measured values were *σ* = 12 spikes/s, *s* in the range 0-50 spikes/s, and *R*_max_ ~ 150 spikes/s (Private Communication with authors of [56] correcting a typographical error in the printed paper). The authors of [56] used mixtures of only two odors at low concentrations in order to avoid widespread activation of ORNs, and hence the measured values of *s* in their experiments are likely low, suggesting stronger suppression in natural conditions (Private Communiation with authors of [56]).

